# Evidence for interacting but decoupled controls of decisions and movements in non-human primates

**DOI:** 10.1101/2024.01.29.577721

**Authors:** Clara Saleri, David Thura

## Abstract

Many recent studies indicate that control of decisions and actions is integrated during interactive behavior. Among these, several carried out in humans and monkeys conclude that there is a co-regulation of choices and movements. Another perspective, based on human data only, proposes a decoupled control of decision duration and movement speed, allowing for instance to trade decision duration for movement duration when time pressure increases. Crucially, it is not currently known whether this ability to flexibly dissociate decision duration from movement speed is specific to humans, whether it can vary depending on the context in which a task is performed, and whether it is stable over time. These are important questions to address, especially to rely on monkey electrophysiology to infer the neural mechanisms of decision-action coordination in humans. To do so, we trained two macaque monkeys in a perceptual decision-making task and analyzed data collected over multiple behavioral sessions. Our findings reveal a strong and complex relationship between decision duration and movement vigor. Decision duration and action duration can co-vary but also “compensate” each other. Such integrated but decoupled control of decisions and actions aligns with recent studies in humans, validating the monkey model in electrophysiology as a means of inferring neural mechanisms in humans. Crucially, we demonstrate for the first time that this control can evolve with experience, in an adapted manner. Together, the present findings contribute to deepening our understanding of the integrated control of decisions and actions during interactive behavior.

**New & noteworthy:** The mechanism by which the integrated control of decisions and actions occurs, coupled or interactive but decoupled, is debated. In the present study, we show in monkeys that decisions and actions influence each other in a decoupled way. For the first time, we also demonstrate that this control can evolve depending the subject’s experience, allowing to trade movement time for decision time and limit the temporal discounting of reward value.

## Introduction

Recent theories and studies argue for an integrated control of decisions and actions during interactive behavior, at least when decisions are rapidly expressed through movements (1–4). According to this framework, decisions, or factors guiding choices such as reward, time and effort, influence action parameters (5–11), and conversely, actions influence decisions (12– 15), even if the choice is about a sensory stimulus or reward harvest during foraging (16–21).

These observations are not compatible with a strict serial organization of behavior, where cognition, including decision-making is separate from perception and action, and lie between the two (22). Instead, they support a dynamic control system operating in a closed loop where movements shape current and future potential actions (i.e. affordances), assigning various importance to the stimuli given such opportunities, and thus directly influencing decisions (23–27).

While the integrated control of decisions and actions is an increasingly established hypothesis, a lively ongoing debate concerns the mechanism by which this integrated control occurs. On the one hand, proposals have suggested that decisions and actions are jointly regulated, possibly by a common source (6, 9, 28, 29). Such “co-regulation” hypothesis, demonstrated in both humans and monkeys, is compatible with the older observations that reward and effort invigorate and slow down both decisions and actions, respectively (8, 30– 34). It has been proposed that a regulation signal computed in the basal ganglia allows to maximize the rate of correct responses by jointly adjusting decisions urgency and action vigor during tasks involving multiple successive choices between actions (6, 35–37).

A second line of research suggests however that decision duration and movement speed are controlled independently of each other, allowing for instance to trade decision time for movement time (19, 20, 38, 39). In support of this possibility, we recently showed that human subjects can shorten their decisions when an accurate and time-consuming movement is requirement (19, 20, 38). We proposed that such adjustment is a “compensatory” mechanism established to decrease the temporal discounting of a positive outcome in each trial, and/or to limit a drop of success rate on a more global time scale. This ability to exchange decision duration for movement duration, i.e. the flexibility to dissociate decision urgency from movement vigor, argues for interacting but decoupled decision and motor control systems.

Importantly, it is not currently known whether this ability to flexibly dissociate decision duration from movement speed is specific to humans, maybe made possible by a sophistication of decision and motor neural circuits through evolution, or whether non-human primates like old-world monkeys, who diverged from the human lineage about 25 MA ago, already have this skill. Likewise, it is currently not known if the regulation mechanism of decisions and actions is exclusive and stable over time or whether it can vary depending on the context in which a task is performed, and whether or not it can evolve with practice.

These are crucial questions to address because a significant part of what we know about the neural substrates of decision-making and motor control comes from (non-human) animal and, more specifically, monkey neurophysiology (40, 41). It is thus important to figure out whether or not the mechanism underlying the integrated control of decision and movement (i.e. joint or compensatory) is similar in both species, stable across contexts (for example, by prioritizing either speed or precision of behavior) and over time, first at the behavioral level in order to then rely on monkey electrophysiology to infer the neural mechanisms in humans.

In the present study, we trained two macaque monkeys to perform a decision between actions task and analyzed decision and movement data collected over dozens of sessions. We focused our analyses on the relationship between decision duration and movement vigor (speed and/or duration), both at the single trial level and between blocks of trials in which motor constraints were varied. We also investigated these relationships as a function of animals’ experience in the task. The analyses strongly indicate that decisions and actions influence each other, but in a decoupled way: decision duration and action vigor could co-vary but also “compensate” each other. Crucially, this integrated control can evolve with practice in an adapted manner. Our data indeed suggest that monkeys can trade movement time for decision time, and vice versa, when the total duration of behavior increases, possibly allowing to limit a drop of reward value due to temporal discounting.

## Methods

### Subjects and ethics statement

The present report describes data collected from two rhesus macaque monkeys (Macaca *mulatta* - monkey G, female, 8 kg, 10 years old, right-handed; monkey B, male, 5 kg, 4 years old, left-handed). Ethical permission was provided by “Comité d’Éthique Lyonnais pour les Neurosciences Expérimentales” (CELYNE), C2EA #42, ref: C2EA42-11-11-0402-004. Monkey housing and care was in accordance with European Community Council Directive (2010) and the Weatherall report, “The use of non-human primates in research.” Laboratory authorization was provided by the “Préfet de la Région Rhône-Alpes” and the “Directeur départemental de la protection des populations” under Approval Number: D69 029 06 01.

### Experimental apparatus

The monkeys sat on a primate chair and made planar reaching movements using a lever held in their dominant hand (Fig. 1A). A digitizing tablet (GTCO CalComp) continuously recorded the lever horizontal and vertical positions (∼100 Hz with 0.013-cm accuracy). Target stimuli and cursor feedback were projected by a VIEWPixx monitor (VPixx Technologies, 120 Hz refresh rate) onto a half-silvered mirror suspended 25 cm above and parallel to the digitizer plane, creating the illusion that targets floated on the plane of the tablet.

**Figure 1.**
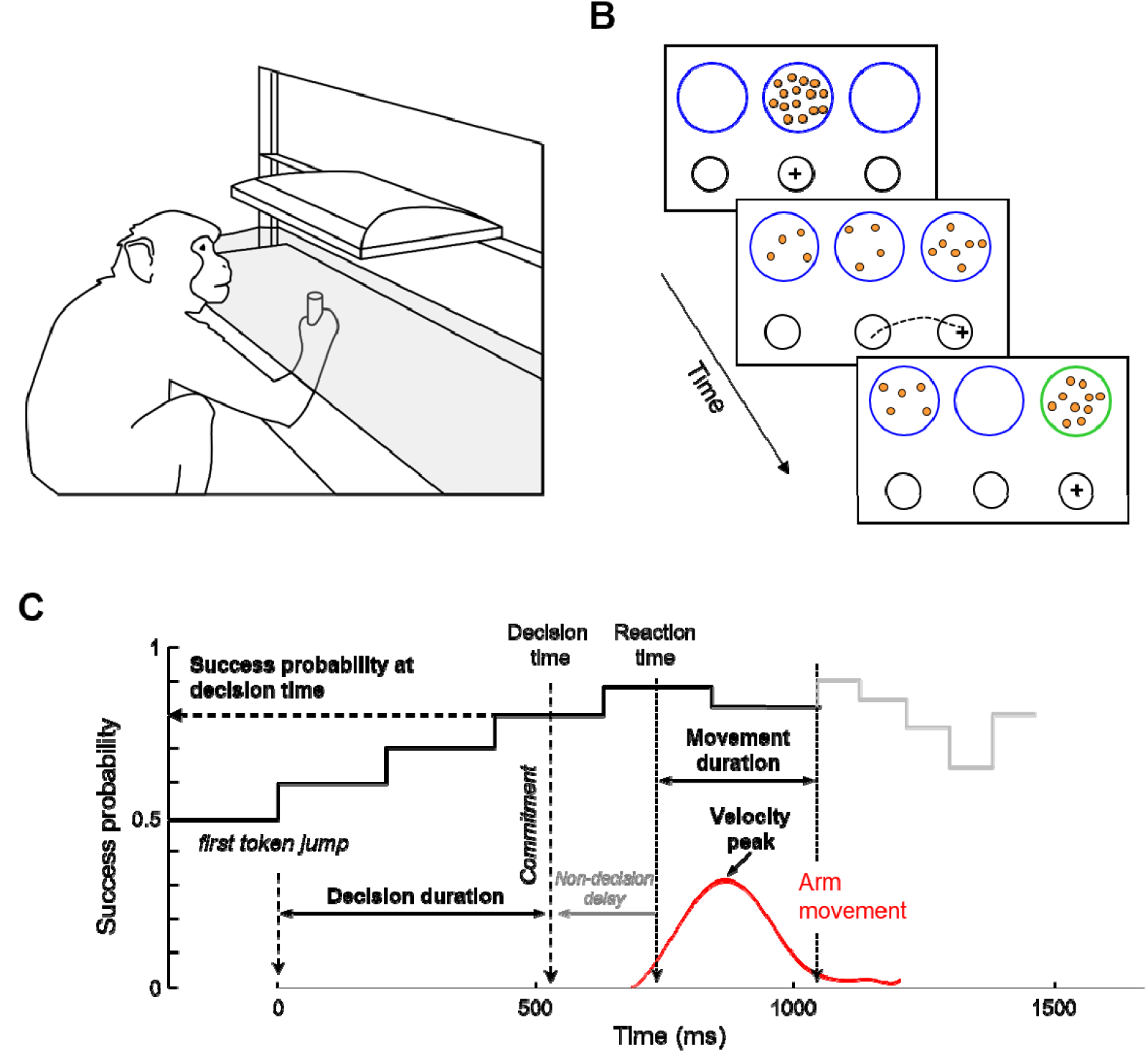
Experimental setup, task and data analysis. *A* - Experimental apparatus. *B* - Time course of a trial in the task. *C* - Temporal profile of success probability in one example trial of the choice task. At the beginning of the trial, each target has the same success probability (0.5). When the first token jumps into one of the two potential targets (the most leftward vertical dotted line), success probability of that target increases to ∼0.6. Success probability then evolves with every jump. Subjects execute a reaching movement (red trace) to report their choices. Movement onset and offset times are used to compute movement duration. Peak velocity is determined as the maximum value between these two events. Movement offset marks the moment when the tokens that remain in the central decision circle jump more quickly to their assigned target (gray trace). The estimated time of the decision is computed by subtracting the subject’s mean non-decision delay estimated in a simple delayed reaching task from movement onset time, allowing computation of the choice success probability at that moment. Only 10 out of 15 jumps are illustrated on this success probability profile.

### Behavioral tasks

The monkeys performed the same task, a modified version of the “tokens task” (see 42 for the original version), as that performed by human subjects and for which data were reported recently (19) (Fig. 1B). They were faced with a visual display consisting of three blue circles (radius: 2.25 cm for monkey G; 2 cm for monkey B) placed horizontally at a distance of 6.375 cm (monkey G) or 7 cm (monkey B) from each other (the “decision” circles). In the central blue circle, 15 small tokens were randomly arranged. Positioned 5 cm (monkey G) or 5.5 cm (monkey B) below, three black circles, organized horizontally as well, defined the “movement” targets. The central black circle, the starting circle, remained constant at a radius of 1.5 cm, while the lateral black circles, positioned at a distance of 8 cm (monkey G) or 7 cm (monkey B) from the center, varied in size (radius: 1.25 -1.75 cm; and 2.25 cm) in separate blocks of trials, resulting in two motor conditions (“small” vs. “large” targets).

A trial started when the animal moved and held the cursor in the black central circle for 500ms. At this point, the tokens began to jump one by one, every 200ms, from the central blue circle into one of the two possible lateral blue circles. The monkeys had to determine which of the two lateral blue circles would contain the majority of tokens before they had all jumped. They reported their decisions by moving the cursor into the corresponding lateral movement target (Fig. 1B). Importantly, monkeys could make and report their choices at any time between the first and last token jumps. However, the movement duration could not exceed 800ms, regardless of the motor condition.

The feedback about timing provided to the monkey was determined online: LabVIEW logged timestamps when the monkey left the start position and when it entered the chosen target, then compared this duration to the maximum allowed duration (800 ms) to ensure the movement was not too long. This differs from the offline analysis, which uses velocity thresholds to determine the onset and offset of movements (see below). If the movement exceeded 800ms (too slow) or reached the target but failed to stop within 800ms (inaccurate), the trial was considered as a movement error and the movement target turned orange. When the choice was properly reported, the remaining tokens jumped more quickly to their final circles (Fig. 1C). This post-decision interval was set to 100-150ms and this acceleration of the remaining tokens implicitly encouraged animals to decide before all tokens had jumped into their respective lateral circles, aiming to save time and increase their reward rate. Note that this feature entails that movement duration carries a temporal cost with respect to the monkey’s rate of correct decisions because the remaining tokens accelerate at movement offset. The visual feedback regarding trial success (both correct decision and correct movement) or failure (correct movement but wrong decision) was provided after the last token jump, with the chosen decision circle turning green or red, respectively. A reward (drops of fruit juice) was given to the monkey after each correct trial performed. Using a pressure dispenser (Crist Instrument) with 35 ms opening duration and 2.0 PSI pressure, 100 rewards equated to 15 ml of juice, corresponding to approximately 0.15 ml per drop. Therefore, monkeys received between 2 and 3 drops of juice per correct trial, depending on their motivation, amounting to approximately 0.30 ml to 0.45 ml of juice. Finally, the next trial began after a 1500ms intertrial interval.

To ensure that the difficulty of decisions was homogeneous among experimental conditions, especially between the two motor conditions, we controlled the sequence of trials experienced by animals in each session. We interspersed among fully random trials (30% of the trials in which each token is 50% likely to jump into the right or the left lateral circle) three special types of trials characterized by particular patterns of token jumps. Between 30 and 40% of trials were so-called “easy” trials, in which tokens tended to move consistently toward one of the circles. Around 20% of trials were “ambiguous,” in which the initial token movements were balanced, making the success probability of each target close to 0.5 until later in the trial. The last special trial type was called “misleading” trials (7-20%) in which the two to three first tokens jumped into the incorrect circle and the remaining ones into the correct circle (please refer to Fig. S1A for a complete description of trial types criteria). In all cases, even when the temporal profile of token jumps of a trial was predesigned, the actual correct target was randomly selected on each trial.

In addition to the tokens task, each animal had to complete trials in a delayed reaching task in each session. The delayed reaching task was similar to the tokens task, but with only one lateral decision circle displayed at the beginning of each trial (on the right or left side of the central circle with a 50% probability). The tokens all moved at the same time (go signal) from the central circle to this unique lateral circle after a variable delay (400-550ms). In this task the reward was given to the monkey if the movement is initiated after the go signal and is accurately executed (endpoint within the target and duration under 800 ms). The target turns orange if the movement is inaccurate or too long. Animals performed this task in the two different motor conditions (small and large targets). The delayed reaching task was employed to estimate the non-decision delay (i.e. the sum of the delays attributable to sensory processing of the stimulus display) as well as to response initiation in each motor condition (Fig. 1C).

### Procedure

Sessions included in the present report where those during which monkeys were proficient in the task, typically following several months of training (see the stability of monkeys’ performance in Fig. 2A). Data included in the analyses were collected over a total of 71 sessions for monkey G and 46 for monkey B. Each session consisted of alternating blocks (between 1 and 5) of 10-60 correct trials in each motor condition whose order alternated on a daily basis. Importantly, the mean success probability profiles of trials in each motor condition were similar (Fig. S1B), ensuring that a potential difference in decision strategy is not due to a difference in the overall difficulty of the trials encountered by each monkey. Monkeys had also to complete between 45 and 60 correct trials of the delayed reaching task in each condition. Then, each session had a predefined goal for the number of correct trials to perform in the main task (often 180 or 240 for monkey G and 240 for monkey B). Depending on the motivation of the animal on a given day, the goal could be adjusted, being either increased or decreased (ranging from 60-240 trials for monkey G and 180-400 for monkey B). Sessions were stopped if cooperation was poor. On average, monkey G performed 174 ± 62 correct trials (both correct decisions and correct movements) by session while monkey B made 253 ± 43 trials.

**Figure 2.**
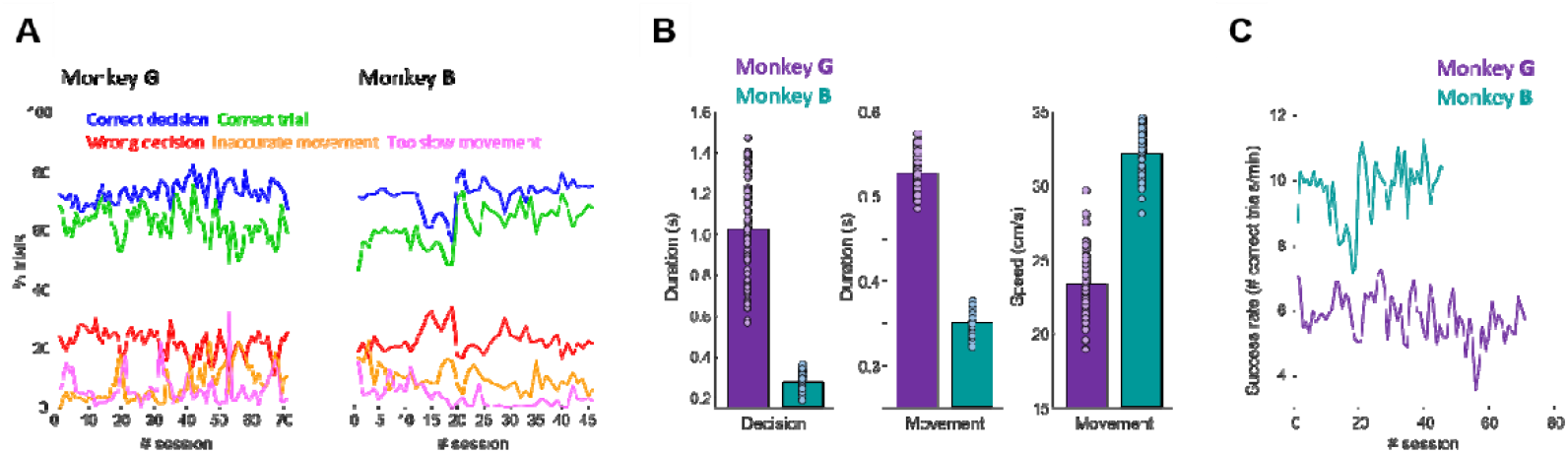
Global performance, decisional and movement vigor in the task. *A* - Performance of each monkey across sessions. *B* - Decision duration, reaching movement duration and speed for each of the two monkeys. Bars illustrate mean values across sessions, and dots illustrate individual values for each session. *C* – Success rate of each monkey across sessions.

### Data analysis

All arm movement data were analyzed off-line using MATLAB (MathWorks). Reaching characteristics were assessed using the animals’ movement kinematics. Horizontal and vertical position data were first filtered using a tenth-degree polynomial filter and then differentiated to obtain a velocity profile. Onset and offset of movements were determined using a 3.75 cm/s velocity threshold. Peak velocity was determined as the maximum value between these two events. To estimate the time at which monkeys committed to their choice (decision timeon each trial, we detected the time of movement onset, defining the animal’s reaction time (RT), and subtracted from it her/his mean non decision delays estimated based on her/his reaction times in the same motor condition (large or small targets) of the delayed reaching task performed the same day. Decision duration was computed as the duration between the decision time and the first token jump (Fig. 1C). The task design allows to calculate, at each moment in time during a trial, the success probability p*i*(t) associated with choosing each target *i* (Eq. 1). For instance, for a total of 15 tokens, if at a particular moment in time the right target contains N_R_ tokens, whereas the left target contains N_L_ tokens, and there are N_C_ tokens remaining in the center, then the probability that the target on the right will ultimately be the correct one, i.e., the success probability of guessing right is as follows:

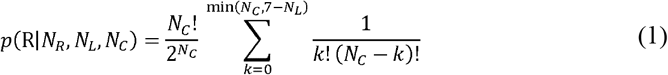

The rate of correct trials was computed for each session as the number of correct trials to complete in that given session divided by the sum of the durations of all trials performed to reach this requested number of correct trials.

### Statistics

Wilcoxon rank sum tests are used to assess the difference in decision duration between categories of trials (e.g. easy versus ambiguous decisions, decisions made in early versus late sessions, etc.). Pearson linear correlations are computed to assess the relationship between decision duration and movement parameters (speed and duration) at the single trial level and between sessions. Pearson linear correlations are also computed the assess the relationship between the degree of coordination between decision duration and movement speed and the duration of decisions between sessions. For all statistical tests, the significance level is set a 0.05.

## Results

### General observations

Over the 71 sessions performed by monkey G, 63 ± 6% of the trials (mean ± standard deviation between the sessions) were successful (both correct choices and correct movements). Without taking into account the success of the movements, 74 ± 4% of her decisions were correct. Movement errors, too slow or inaccurate, were relatively sparse, representing 7 ± 5% and 8 ± 6% of the total trials, respectively. Monkey B’s performance over the 46 sessions he completed was comparable to that of monkey G. 62 ± 7% of his trials were successful (correct decisions and correct movements). Regardless of movement outcome, 73 ± 5% of his choices were correct. 4 ± 4% of his movements were too slow, 10 ± 4% were inaccurate. For both monkeys, performance was stable across the sessions included in the present report (Fig. 2A).

Although their performance in the task is comparable, the analysis of animals’ duration of decisions, duration and speed of arm movements suggests that they used different strategies to achieve these similar outcomes (Fig. 2B). On average, the duration of monkey G’s decision was 1026 ± 242ms. As expected, she decided more quickly when trials were easy compared to when they were ambiguous (872 ± 232ms vs. 1154 ± 259ms, Wilcoxon rank sum test, z = -5.7, p<0.0001). To report her decisions, monkey G made arm movements whose duration was 527 ± 21ms and their speed was 23 ± 2 cm/s. Monkey B was on average much faster to decide and act compared to monkey G. His decision duration was 249 ± 43ms, with faster choices when trials were easy compared to when they were ambiguous (230 ± 40ms vs. 369 ± 73ms, Wilcoxon rank sum test, z = -6.6, p<0.0001). To report his decisions, monkey B made movements whose speed was 32 ± 3 cm/s and their duration was351 ± 14ms. As a consequence, the rate of correct trials, defined for each session as the number of correct trials per minute (Fig. 2C), was much higher in monkey B than in monkey G (9.7 ± 1 correct trials/min vs. 5.8 ± 0.7 correct trials/min).

### Relationship between decision duration and movement vigor at the single trial level

To study the relationship between decision duration and arm movement vigor (speed and duration) *within trials*, we first grouped all trials (correct and error trials, regardless of motor condition) across sessions, separately for each monkey, according to decision time computed in bins of 200ms, and calculated the mean (± standard error) values of movement parameters in each bin.

The analysis of monkey G movement speed as a function of decision duration (including decision durations until 2200ms, which corresponds to 99.6% of the trials) shows a slight but continuous *increase* of movement speed with decision duration (from 22.5 to 23.9 cm/s, Fig. 3A, left panel). A linear regression through the data indicates a very strong and significant relationship between the two variables (Pearson correlation, r=0.98, p<0.0001), with a positive slope (0.14) and an intercept at 22.4 cm/s. No significant relationship between decision duration and movement duration is observed (Pearson correlation, r=0.02, p=0.9).

**Figure 3.**
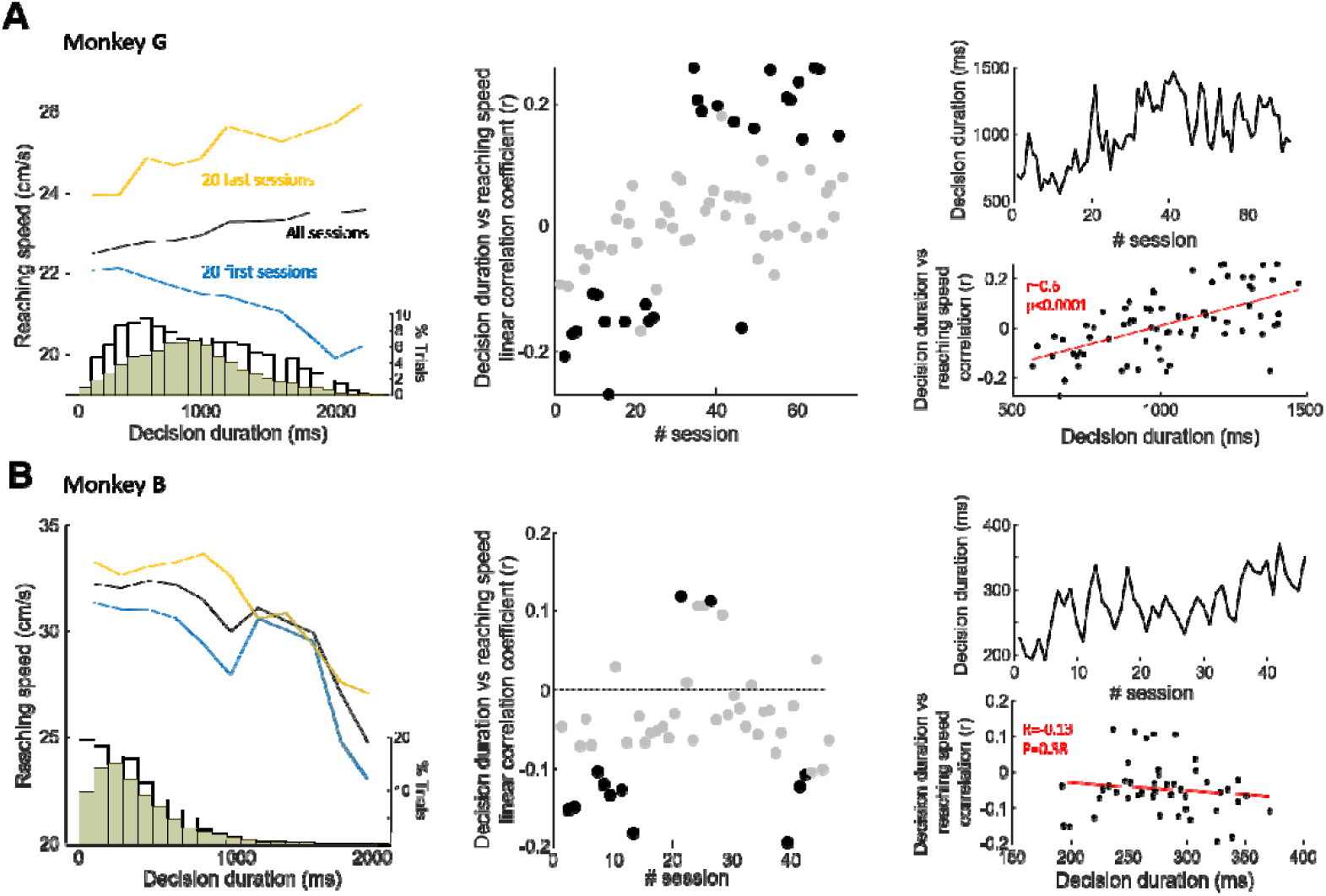
Relation between decision duration and arm movement speed within trials. *A – Left panel*: Speed of monkey G’s arm movements (reaching) as a function of decision duration, computed in bins of 200ms, averaged (± standard error, shaded areas) across all trials (black), trials of the 20 first sessions (“early” sessions, blue) and trials of the 20 last sessions (“late” sessions, orange). Histograms at the bottom of the panel show the distributions of the duration of monkey G’s decisions in the two groups, “early” and “late” sessions. *Center panel*: Pearson linear correlation between the duration of monkey G’s decisions and arm movement speed (velocity peak) as a function of the session performed by the monkey. Each dot illustrates the value for a given session. The blue and orange shaded areas highlight the 20 first and the 20 last sessions performed by monkey G, respectively. A negative (positive) r value means that the longest the decision, the slowest (fastest) the movement. The black filled dots mark sessions in which the correlation is significant. *Right panel, top*: Mean duration of monkey G’s decisions within sessions, as a function of the session performed by the monkey. *Right panel, bottom*: Relationship between the Pearson correlation between decision durations and the speed of monkey G’s movements and the mean duration of her decisions in each session. Each dot illustrates the value for a given session. The red line illustrates the result of a linear regression through the data. *B* – Same as in A for trials performed by monkey B.

By contrast, the analysis of the relationship between the duration of decisions and the speed of monkey B’s movement within trials (including decision durations until 2200ms, which corresponds to 99.9% of the trials) shows a continuous *decrease* of movement speed with decision duration (from 32.2 to 25 cm/s, Fig. 3B, left panel). A linear regression through the data indicates a significant relationship between the two variables (Pearson correlation, r=-0.85, p=0.0008), with a negative slope (-0.62) and an intercept at 34 cm/s. As for monkey G, no significant relationship between monkey B decision duration and movement duration is observed (Pearson correlation, r=-0.05, p=0.9).

We then asked whether this relationship between decision duration and movement speed within trials was stable over extended periods of time, depending on the monkeys’ experience in the task, or if it could possibly evolve depending on this experience. To do this, we first carried out the same analyzes as those described above but with trials grouped according to the number of sessions performed by each monkey. For both animals, we compared trials collected during their first 20 sessions (“early” sessions) with those performed in their last 20 sessions (“late” sessions).

The relationship between the duration of decisions and the speed of monkey G’s movements is strong and significant in both groups of trials (“early” sessions: r=-0.9, p=0.001; “late” sessions: r=0.82, p=0.0021), but the slope is *negative* for “early” sessions (-0.22) and *positive* for “late” sessions (0.19). This means that the longest decisions were followed by the slowest movements during monkey G’s first sessions and then, with practice, this relationship reversed (Fig. 3A, left panel; Fig. S2A). To better capture this evolution, we analyzed the session-by-session relationship between decision duration and action speed in monkey G and we found that a linear regression shows a strong and significant fit (Pearson correlation, r=0.61, p<0.0001), with a positive slope, meaning that the link between decision duration and movement speed continuously evolved through monkey G’s practice (Fig. 3A, middle panel). In particular, among the first 20 sessions, decision durations were significantly correlated with movement speed in 8 sessions, always with long decisions followed by slow movements. Among the last 20 sessions, we found a significant correlation between decision duration and movement speed in 8 sessions too, but this time always with long decisions followed by rapid movements. Finally, we analyzed the duration of monkey G’s decision in the two groups of trials and noticed that with practice, she became more conservative and decided more slowly (mean ± standard error; “early” sessions: 760 ± 6ms vs. “late” sessions: 1101 ± 7ms, Wilcoxon rank sum test, z = -35, p<0.0001) (Fig. 3A, left panel). Interestingly, we found a positive and significant correlation between the degree of coordination between decision duration and movement speed and the duration of decisions between sessions (Pearson correlation, r=0.6, p<0.001) (Fig. 3A, right panels). In other words, the emergence of the positive correlation between decision duration and movement speed (the longest the decision, the fastest the movement) within trials through practice coincides with the increase of the duration of monkey G’s decisions through practice. In the supplementary section of the present report, we describe the relationship between the evolution of the correlation of decision duration and movement speed and the evolution of decision duration in two other monkeys who performed a similar version of the task (Fig. S3), strongly supporting the same conclusion (Fig. S4).

As for monkey G, the relationship between the duration of decisions and the speed of monkey B’s movements is significant in both groups of trials (“early” sessions: r=-0.78, p=0.0044; “late” sessions: r=-0.9, p=0.0002). But contrary to monkey G, the slope is negative in both groups: -0.65 for “early” and -0.64 for “late” sessions (Fig. 3B, left panel; Fig. S2B). This means that regardless of the monkey’s level of experience in the task, the longest decisions were followed by the slowest movements. We did not find a particular link between the relationship between decision duration and action speed and the sessions performed by monkey B through his practice (Fig. 3B, middle panel). Among the first 20 sessions, decision durations were significantly correlated with movement speed in 7 sessions, and always with long decisions followed by slow movements. Among the last 20 sessions, the correlation between decision duration and movement speed was significant in 3 sessions, still with long decisions followed by rapid movements. Interestingly though, despite the fact that the relationship between decision duration and movement speed *within trials* did not change with practice, we found that movement speed was overall higher in the “late” sessions group (intercept = 35 cm/s) compared to the “early” sessions one (intercept = 32.9 cm/s) (Fig. 3B, left panel). The analysis of the duration of monkey B’s decision in the two groups showed that he was slightly more conservative in the “late” sessions compared to the “early” sessions (302 ± 3ms vs. 259 ± 3ms, Wilcoxon rank sum test, z = -17, p<0.0001) (Fig. 3B, left panel).

Contrary to monkey G, no significant relationship between the degree of coordination between decision duration and movement speed and the duration of monkey B’s decisions between sessions is observed (Fig. 3B, right panels).

### Effect of motor constraints on the relationship between decision and action

Another way to study the coordination between decision duration and movement vigor is to encourage animals to vary the vigor of their movements in blocks of trials and assess the consequence of this adjustment on decision duration. A co-regulatory mechanism predicts that slowing movements will be accompanied by slowing decisions, while a more flexible mechanism could be demonstrated by observing shortened decisions when movements are longer, as we have seen in humans (19, 20, 38).

Across all sessions, when movements were correctly executed (decisions could be correct or wrong), the speed of monkey G’s movements was slower (mean speed ± standard error: 22.8 ± 0.05 cm/s vs. 23.3 ± 0.05 cm/s, Wilcoxon rank sum test, z = -6, p<0.0001) and their duration longer (536 ± 1ms vs. 525 ± 1ms, z = 8.4, p<0.0001) when they were executed toward the small targets (“small” target block) compared to when they aimed to the large targets (“large” target block). We found that the duration of monkey G’s decisions was overall slower when movements were directed toward small targets compared to when they reached the large targets (1020 ± 6ms vs. 953 ± 6ms, z = 8.6, p<0.0001). At the session level, we found a significant effect of the motor condition on the speed and duration of monkey G’s movements in 16 and 19 out of 71 sessions, respectively (Wilcoxon rank sum test, p<0.05), with movements being faster and shorter when directed to the large targets in 15/16 and 18/19 cases, respectively (Fig. 4A, left and middle panels). The duration of monkey G’s decisions was modulated by the motor condition in 34 out of 71 sessions (Wilcoxon rank sum test, p<0.05), with decisions being slower when movements were slower in the majority of cases (27/34) (Fig. 4A, right panel). Interestingly, we found that this tendency to slow down decisions when a time consuming movement was necessary diminished with practice. Indeed, while the effect of the motor condition on movement speed and duration was stable over monkey G’s practice (Fig. 4A, left and middle panels), the number of sessions during which long decisions preceded long movements decreased (20 over the first 35 sessions; 5 for the last 35 sessions) (Fig. 4A, right panel).

**Figure 4.**
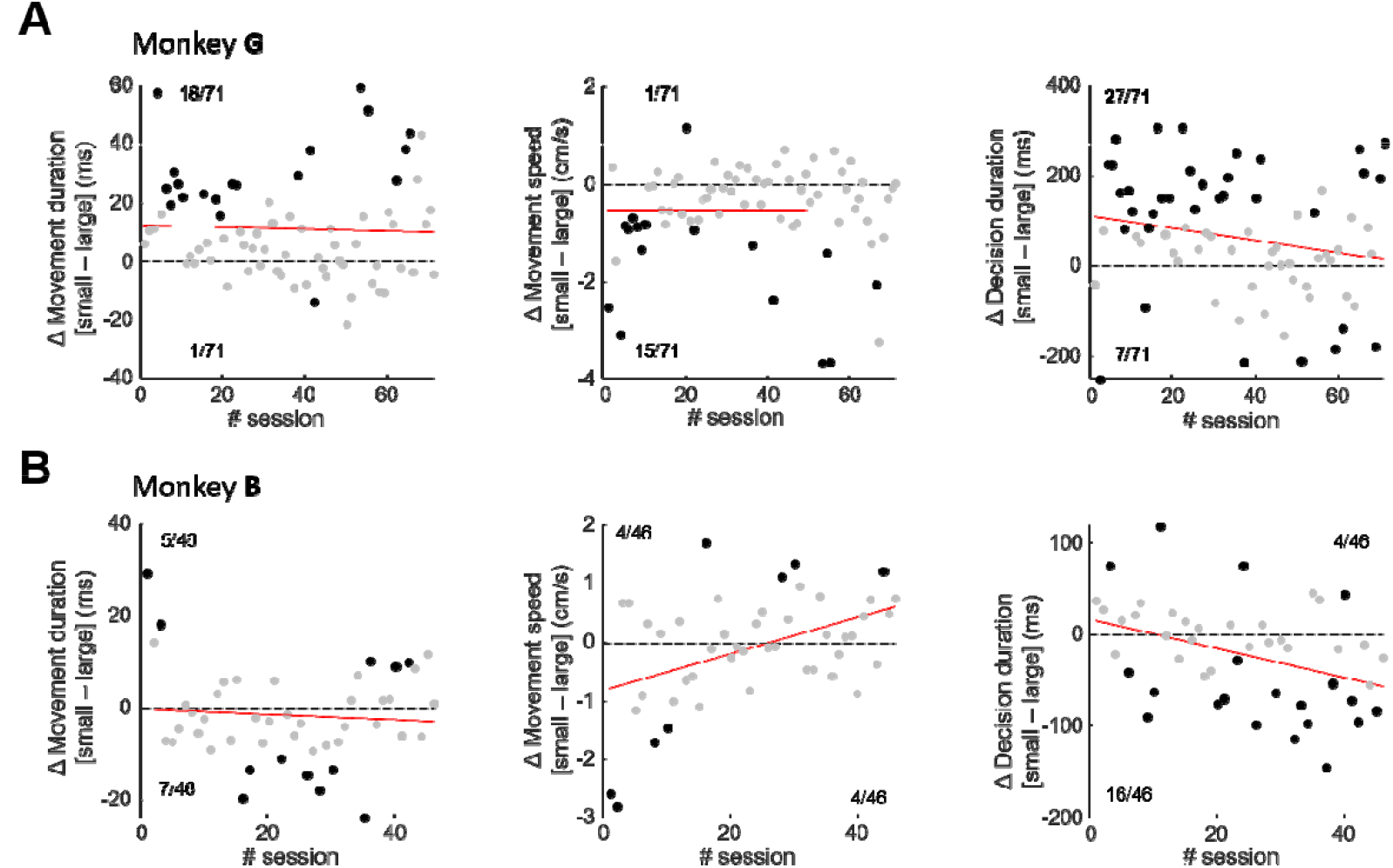
Effect of motor constraints on arm movement vigor and decision duration. *A – Left panel*: Difference of arm movement duration between trials of the small target blocks and those of the large target blocks, as a function of the session performed by monkey G. Each dot illustrates data of a given session, and the black dots mark sessions in which the difference is statistically significant (the number of statistically significant cases is reported for each difference direction). A positive value indicates that movements were longer when they were executed toward the small targets compared to when they were executed toward the large targets. The red line illustrates the result of a linear regression through the data. *Center panel*: Same as in A for the difference of the speed of monkey G’s arm movement between the two motor conditions. *Right panel*: Same as A for the difference of the duration of monkey G’s decisions between the two motor conditions. *B* – Same as in A for trials performed by monkey B.

The effect of target size on the speed and duration of monkey B’s movement is less clear, at least when trials are grouped across sessions. Movement speed was not significantly modulated as a function of the motor condition (small vs. large target: 32 ± 0.07 cm/s vs. 32.2 ± 0.07 cm/s, Wilcoxon rank sum test, z = -0.2, p=0.8). Movement duration was slightly longer when movement reached the large targets compared to when they reached a small target (349 ± 0.7ms vs. 351 ± 0.7ms; z = -2.3, p=0.02). At the session level, the effect of target size on the kinematics of monkey B’s movements (duration and speed) was mixed, as can be seen in figure 4B, left and middle panels. However, target size significantly modulated decision duration in 20 out of 46 sessions (Wilcoxon rank sum test, p<0.05). In the majority of cases (16/20), decision duration was longer when movements reached the large targets (Fig. 4B, right panel). Interestingly, with practice, this monkey progressively decreased his movement speed in one condition (“large” target block) compared to the other one (“small” target block) (Fig. 4B, middle panel), and this relative decrease of speed was accompanied by a session-by-session lengthening of decisions in the large target blocks compared to the small target blocks (Fig. 4B, right panel). The link between these two adjustments from one session to the next is significant (Pearson correlation, r=-0.3, p=0.03).

## Discussion

In the present report, we show in non-human primates that the duration of perceptual decisions and the speed of the arm movements executed to report these choices are often correlated at the single-trial level. The results support an interacting but decoupled control of the decision-making and movement execution processes: decision duration and action speed are sometimes co-regulated (the slower the decision, the slower the movement), and they sometimes “compensate” each other (the slower the decision, the faster the movement). We also demonstrate that the degree and direction of coordination between decision duration and movement vigor can evolve, particularly as a function of the overall duration of the animals’ decisions in each session: the “compensatory” mode of regulation comes into play as duration of choices increases. Finally, we show that when the motor context encourages time-consuming movements, decision duration and movement vigor are often co-regulated. This context-dependent relationship also appears flexible depending on the animals’ experience in the task.

These observations agree with many recent experimental results that challenge the classic view of behavior organization (44), in which perception, decision, and action are considered as independent, temporally separate and serial processes (22). More and more studies indeed show that decisions and actions are closely interconnected and influence each other, at least during rapid interactive behavior (5, 7–21, 28, 29, 38, 45).

A particularly interesting question concerns the mechanism by which such integrated control of decisions and actions occurs. On the one hand, decisions and actions might be coupled, jointly regulated by a common source (6, 9, 28, 29). According to this hypothesis, in a context encouraging decision speed, the fastest choices will be reported with the fastest movements. Similarly, if a decision needs to be reported with a slow and careful movement, this decision will tend to be longer than that expressed by a rapid movement. A second line of research suggests however that the control of decision duration and movement speed is decoupled, allowing for instance to trade decision time for movement time if necessary (19, 20, 38, 39, 46, 47). At the origin of this second hypothesis, we showed that human subjects can shorten their decisions when an accurate and time-consuming movement is required (19, 20, 38). We proposed that such adjustment is a compensatory mode possibly established to prevent the temporal discounting of a positive outcome in each trial, and/or to limit a drop of success rate on a more extended time scale. Interestingly, in two recent studies specifically designed to address the mode of regulation of decisions and actions (28, 29), and whose results support a co-regulation mechanism, decision and movement durations had no impact on the participants’ success rate (i.e. it was not adaptive in terms of success rate to speed up movements after a long decision for instance), unlike the design of the experiments described in Reynaud et al. (19) and Saleri Lunazzi et al. (20, 38).

The data presented in this report suggest for the first time that both modes of coordination can be used by the same subject. At the single trial level, the slowest decisions made by monkey B were expressed with the slowest movements (Fig. 3B, left panel), a result consistent with a co-regulation of decisions and actions. Monkey G also co-regulated but mostly during the first sessions she performed. With experience, the relationship between the duration of her decisions and the speed of her movements reversed, consistent with a decoupled, “compensatory” mode of regulation of choices and actions (Fig. 3A, left panel). With trials grouped as a function of animals’ experience, we found in both monkeys that decisions were slower in the late sessions compared to the early sessions, and these slower decisions were reported with faster movements (Fig. 3A-B, left panels), a pattern of results also consistent with a “compensatory” mode of regulation of decisions and actions. Finally, with trials grouped according to the motor context in which decisions were made (i.e. small vs. large targets), we found evidence for a co-regulation of decisions and actions, with slow movements often preceded by slow decisions, with a trend for an evolution toward a “compensatory” mode in monkey G (Fig. 4A). The way in which the duration of the decision and the vigor of the action are coordinated thus appears complex, flexible and dynamic depending on the context in which the task is carried out (especially the level of experience in the task). In the following paragraph, we discuss the possible reasons which could explain the use of these different modes of coordination.

It is well-established that individuals tend to exhibit a preference for policies that maximize their rate of correct responses when faced with multiple successive decisions (48, 49). Several factors influence this success rate, with time being a crucial one, as evidenced by the well-known concept of temporal discounting of reward value (50). During movement execution for instance, action duration and speed are two (strongly related, Fig. S5) parameters that partially describe a single latent variable, the vigor, that reflects movement’s utility and that the brain seeks to optimize as a function of the economic context (speed-accuracy trade-off, experience in the task, etc.) in which a behavior occurs (4, 34, 51, 52). Because both decisions and actions are time consuming processes, minimizing the time required to obtain a reward likely leads to adaptations concerning both decisions and movements. A decoupled yet interconnected control of decisions and actions provides individuals with the ability to adapt both deliberation and movement speed together, minimizing the time it takes to achieve a given goal. Moreover, this mode also possesses the flexibility to selectively adjust one aspect while preserving the integrity of the other. For instance, during easy decisions, deliberation time can be slightly sacrificed when accurate and slow response movements are required (19, 20, 38). The present results indicate that just like humans, monkeys also have the ability to flexibly adjust the way they coordinate their decisions with their actions, in particular according to their experience in the task.

The evolution of the coordination between decision duration and action vigor depending on animals’ experience is particularly useful in revealing the principles underlying how these two processes are controlled. Crucially, the emergence of the compensatory mode of coordination always coincides with an increase of the duration of the animal’s overall behavior (Fig. 3A, right panel; Fig. S4). This is particularly salient in monkey G who showed an important increase in her decision durations over the course of the sessions carried out (Fig. 3A, right panel). Interestingly, the subtle and gradual decoupling of decision and action observed across sessions between large and small target conditions (Fig. 4A, right panel) occurs simultaneously with the overall lengthening of her decision durations across sessions (Fig. 3A, top right panel). She may have wanted to seek to compensate for her long movements by limiting the increase in the duration of her decisions in the most constrained motor blocks (i.e. small target). By contrast, monkey B, who was constantly very fast to decide and act, showed almost no sign of compensation between decision and movement durations. In monkey S and monkey Z who were even slower to decide than monkey G, the coordination within trials between decision duration and action speed is only of the compensatory type, but it still depends on the duration of the animals’ overall decisions (Fig. S4). We thus propose that the decoupled, compensatory mode of regulation is highly adaptive in the sense that it allows to deal with deadlines and/or makes it possible to limit the temporal discounting of the reward value in each trial. In the longer term, it could also help optimize the overall success rate of the animals in each session. Between these three possibilities, we lean toward the limitation of the temporal discounting of the reward value in each trial because (1) monkeys almost never failed their trials because of responding after 3s (i.e. the deadline of deliberation) and (2) the overall success rate of monkey G does not appear to improve despite the emergence of the compensatory mode of regulation across sessions (Fig. 2C).

The different strategies adopted by the two monkeys can be explained by their very different personalities and backgrounds. Monkey B who faced his first experimental task is much younger than monkey G. He adopted such a fast and impulsive behavior that there was almost no temporal discounting of reward in each (correct) trial, and no room for either decision or movement duration shortening. Consequently, a decoupled mode of regulation that would compensate for slightly longer decisions or movements to meet some reward rate maximization principles was probably irrelevant to this monkey. Monkey G however, who is older and more experienced in laboratory experiments, exhibited a more conservative and less vigorous behavior. Therefore, she had to face higher temporal discounting of reward in each trial and lower reward rate at a more global scale (Fig. 2C). Compensation between decision and movement duration becomes beneficial in this case.

Human subjects having performed the same task show a compensatory mode of coordination of decision and action when the trials are grouped according to the motor constraints imposed on the subjects (19). By contrast, with the same comparison (decision durations between small versus large target conditions), we report here that monkeys tend to primarily exhibit a co-regulation of decisions and actions. This difference between monkeys and human subjects can be explained by at least two factors. Firstly, monkeys received drops of juice at each correct trial as a reward, so their strategy was probably primarily driven by the prospect of receiving this strong primary reinforcer. By contrast, humans did not receive instant rewards. Their motivation was likely more oriented toward the completion of the required number of correct trials as quickly as possible, aiming to maximize their overall success rate at the sessions level. Another, not mutually exclusive explanation is that taking the motor context into account to compensate for movement duration during the decision-making process requires a significant level of abstraction that monkeys can only gradually acquire with training and experience, to eventually exploit it in an adapted manner. Interestingly, we report that with practice, the slowdown of monkey G’s decisions reported with slow movements in specific blocks of trials tends to decrease (Fig. 4A). By mastering the timing parameters of the task over the sessions, she may have realized that beyond accuracy at the single trial level, she could also optimize her time spent in a session by slightly reducing the duration of her decisions when movements were slow, just as human subjects did, with the difference that they were able to do it after just one session.

In previous experiments in which humans and monkeys were tested in a task similar to that described in the present report, we proposed that both decision urgency and movement vigor (6, 9) were co-regulated. This conclusion was based on two observations. At the single trial level first, early decisions (usually made on the basis of strong sensory evidence but low urgency) were followed by long duration movements whereas later decisions (relying on weak sensory evidence but stronger urgency) were followed by shorter and faster movements, as if they were influenced by the strong level of urgency encountered when a significant amount of time has elapsed during a trial. Second, when subjects were encouraged to make earlier and less accurate decisions in blocks of trials (fast speed-accuracy trade-off regime), we observed faster movements compared to a condition where accuracy was emphasized. The multiple modes of regulation of decisions and actions that we report in the present study suggest however that the mechanism underlying the control of decision duration and movement speed is more elaborated than being under the influence of a single regulation signal. Indeed, while a shared signal may control decision and movement durations at the single trial level (especially when fast movements follow long decisions), and between different decisional speed-accuracy trade-off regimes, a unique signal cannot account for the fact that the movements executed following the slower decisions made during the late sessions were faster compared to those made during the early sessions during which decision were faster. Together, these observations suggest the involvement of multiple sources of regulation of decisions and actions, likely implemented by diverse brain circuits, and used differently and specifically depending on the context in which a task takes place (level of experience in the task, speed-accuracy trade-off, etc.).

It is important to note that the present results are all correlational. A next very interesting question concerns the relationship between the average duration of behavior and the decision-action coordination mode used by the subjects. In this report we propose, on the basis of correlational data, that this mode is determined by the duration of decisions but this must be tested directly, if possible in a way that tends toward the establishment of a causal link between the two phenomena. A possibility of intervention concerns electrical microstimulation in the monkey cerebral cortex, especially in areas involved in the determination of choice. For example, we have shown in the past that electrical stimulation in the dorsal part of the premotor cortex and, to a lesser extent, in the primary motor cortex, could lengthen decisions if this stimulation was delivered shortly before choice commitment occurs (53). By experimentally prolonging the duration of decisions, it would then be possible to observe movements executed rapidly in order to compensate for this additional decisional duration.

Electrophysiological data have indeed highlighted an overlap of brain regions involved in decision-making and those of action processes, particularly in the sensorimotor cortex and the basal ganglia (1, 40, 54). Studies suggest, for example, that activity in the internal section of the globus pallidus, the output nucleus of the basal ganglia, determines decision urgency (35, 36) as well as movement vigor (55–57). In the present study, we describe a within trial level of coordination between decisions and actions possibly established to deal with deadlines and promote the value of the immediate reward. This mechanism might be implemented within the cortico-basal ganglia-thalamo-cortex circuits, with the dopamine as a transmitter dedicated to the exploitation of the local reward opportunities (58). Interestingly, a recent study conducted by Herz and colleagues (39) shows that the subthalamic nucleus (STN) can independently control movement and decision speed in distinct time windows, a result that is compatible with the contribution of this basal ganglia nucleus to the decoupled coordination of decisions and actions that we report at the behavioral level in the present study. We also propose that the different modes of coordination of decisions and actions may serve more global, abstract goals such as the optimization of the rate of successes at the session level. This mechanism might be implemented in broader circuits under the regulation of neuromodulation systems, such as the noradrenergic circuits (59–61). Another possibility, not mutually exclusive, is that structures known to perform predictive computing and build internal models of the world, such as the cerebellum, would play a role in such long term goal establishment (26, 62), by building internal models of behavior utility in each task, allowing to optimize the global value of behavior over extended time scales.

## Supplemental material

Supplemental Figs. S1-S5: https://doi.org/10.6084/m9.figshare.27125388

