## Supplemental material for "Evidence for interacting but decoupled controls of decisions and movements in non-human primates"

*Running head*:

Integrated control of decisions and movements in non-human primates

*Corresponding author information:*

David Thura

Lyon Neuroscience Research Center – Impact team

INSERM U1028 – CNRS UMR5292 – University Claude Bernard Lyon 1

16 avenue du Doyen Jean Lépine, 69675 Bron, France

### Supplementary information

Relation between success probability and trial duration as a function trial types and motor conditions

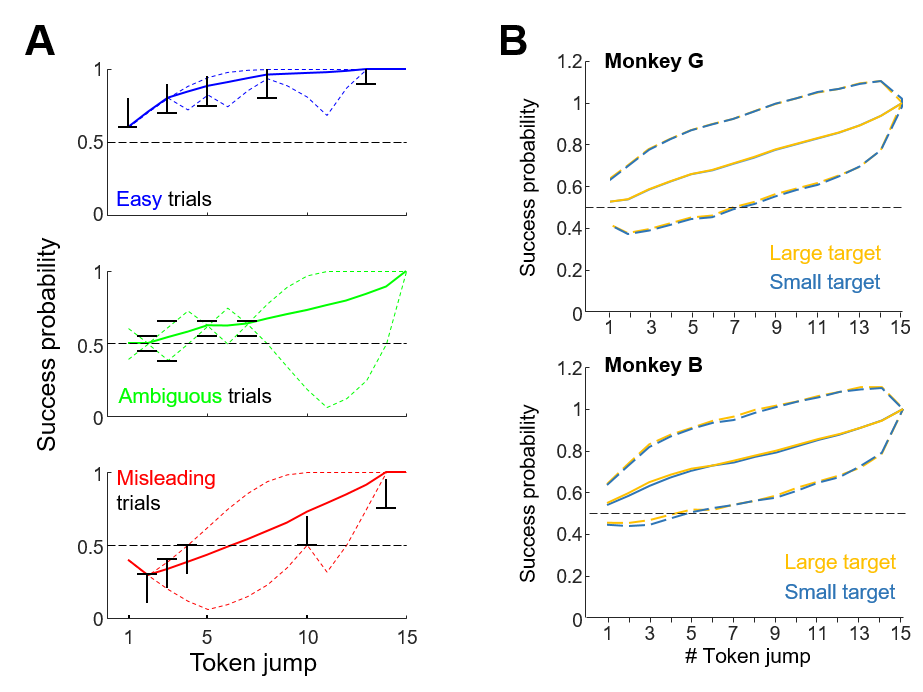

**Figure S1**: A - Average (solid curves) and range (dotted curves) of success probability (SP) profiles of trials classified as “easy” (blue), “ambiguous” (green), or “misleading”. To be classified as an easy trial, the SP profile of the trial with respect to the correct target had to meet the following criteria (black marks): >0.6 after the 1st token jump, >0.7 after 3 jumps, >0.8 after 5 jumps, >0.8 after 8 jumps, >0.9 after 13 jumps. The criteria for an ambiguous trial were the following: SP = 0.5 after 2 jumps, between 0.38 and 0.65 after 3 jumps, between 0.55 and 0.65 after 5 jumps and after 7 jumps. Finally, the criteria for “misleading” trials were: SP <0.3 after 2 jumps, <0.4 after 3 jumps, <0.5 after 4 jumps, >0.5 after 10 jumps, and >0.75 after 14 jumps. The SP profile of the other trials (30%) could take any shape except those corresponding to the special trial types mentioned above. B – Average (± SD) success probability profiles of all trials, with respect to the correct target, experienced by monkey G (top) and monkey B (bottom) during blocks of trials where targets were large or small.

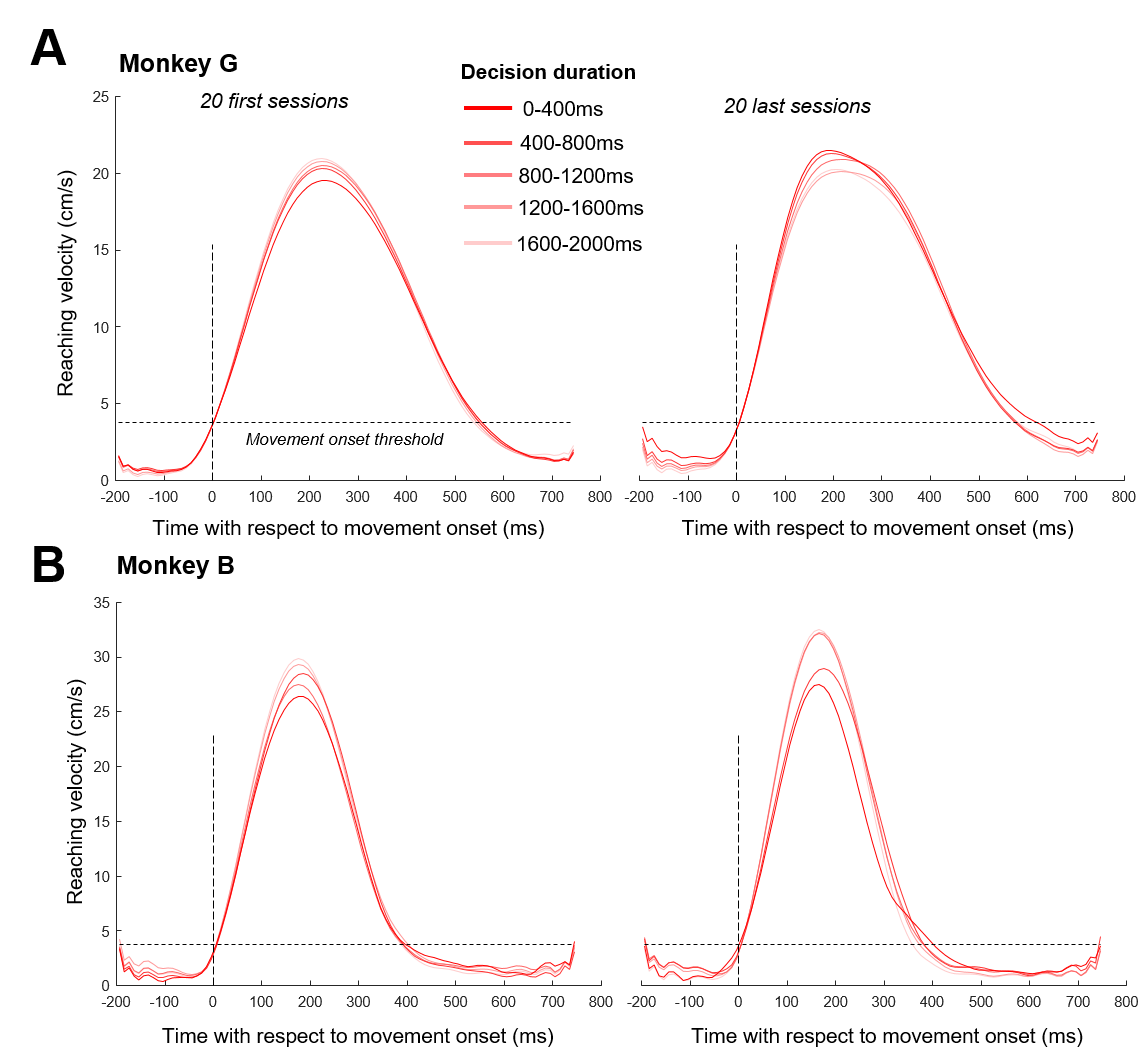

**Figure 2: Velocity profiles of reaching movements executed by monkey G (A) and monkey B (B) as a function of their decision duration and their experience in the task.** Trials are grouped as a function of decision duration binned every 400ms, from 0ms to 2000ms, for the 20 first (left panels) or for the 20 last sessions performed by monkeys.

In the supplementary information section of the present report, we describe a subset of data collected on two rhesus macaque monkeys (Macaca *mulatta* - monkey S, male, 7 kg, 6-8 years old; and monkey Z, male, 5 kg, 4-6 years old) trained and tested by DT in his previous affiliation (Université de Montréal, Montréal, QC, Canada), in the laboratory of Paul Cisek. Several reports involving these two monkeys have been published, including details of the ethical and institutional permissions for these experiments (1–4).

These two monkeys performed the original version of the “tokens” task (5). It is the same task as that performed by monkey G and monkey B, except that in this version, only decisions circles were presented to the animals. The decision rule by itself was exactly the same as that experienced by monkey G and monkey B. Monkeys had to guess which of the two lateral circles would receive the majority of the jumping tokens at the end of the trial. To report their choices, monkeys moved a lever with their hand from the central circle (in which the 15 tokens were randomly located at the beginning of each trial) to one of the two lateral decision circles (Fig. S3).

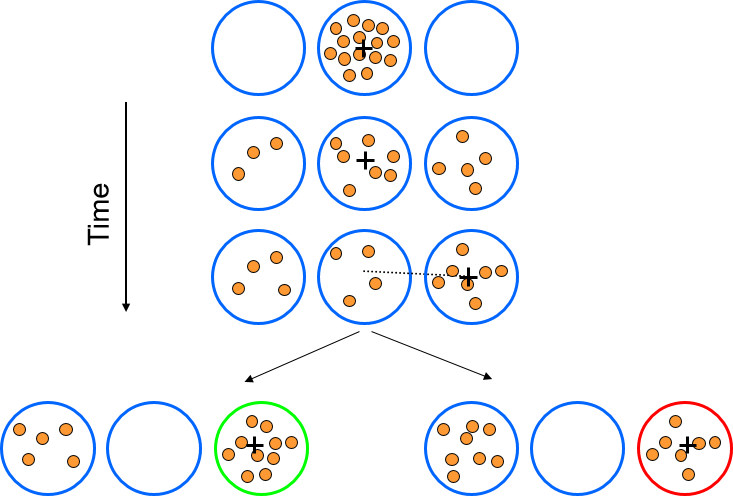

**Figure S3 - The "tokens" task performed by monkey S and monkey Z.** During each trial of the “tokens task”, 15 tokens jump, one every 200ms, from the central circle to one of two outer target circles. The monkey’s task is to move the lever (identified on the screen with a black cross) to the target that will ultimately receive the majority of the tokens.

As monkey G and monkey B, monkey S and monkey Z performed between 30 and 60 successful trials of a delayed reach (DR) task in each session.

In the following paragraphs we report the information which seems essential to us to understand the results in the context of one of the questions addressed in the main manuscript, namely the coordination of the duration of decisions with the vigor of movements at the single trial level, the evolution of this coordination according to the sessions carried out by the animals, and the link between the duration of decisions and the level of coordination between decisions and actions. The complete methodological information about experiments in which monkey S and monkey Z were involved is described in (4).

Data include 261 sessions performed by monkey S (129595 trials; mean number of trials per session ± SD: 497 ± 250) and 279 sessions performed by monkey Z (146756 trials, 526 ± 175 trials per session). The sessions could have been performed when monkeys’ head was free to move during behavioral sessions, and when it was fixed during behavioral sessions only or during electrophysiological sessions.

In this supplemental section we focus our analyses on the duration of monkeys’ decisions as well as the speed (i.e. the peak of velocity) of their movements. All arm movement data were analyzed exactly the same way as in the main manuscript, i.e. off-line using MATLAB (MathWorks). Horizontal and vertical position data were first filtered using a tenth-degree polynomial filter and then differentiated to obtain a velocity profile. Onset and offset of movements were determined using a 3.75 cm/s velocity threshold. Peak velocity (VP) was determined as the maximum value between these two events. To estimate the time at which monkeys committed to their choice (decision time, DT) on each trial, we detected the time of movement onset, defining the animal’s reaction time (RT), and subtracted from it her/his mean non decision delays (ND) estimated based on her/his reaction times in the delayed reach task performed the same day. Decision duration (DD) was computed as the duration between DT and the first token jump.

Over the 261 sessions performed by monkey S, 77 ± 3% of the trials (mean ± standard deviation between the sessions) were successful (both correct choices and correct movements). Because target size was large (Ø = 3.5cm), movement errors were extremely rare. On average, the duration of monkey S’s decision was 1232 ± 192ms, and the speed of his movements was 23.5 ± 3.1 cm/s. Over the 279 sessions performed by monkey Z, 75 ± 3% of the trials (mean ± standard deviation between the sessions) were successful (both correct choices and correct movements). As for monkey S, because target size was large, movement errors were extremely rare. On average, the duration of monkey S’s decision was 1145 ± 260ms, and the speed of his movements was 24.7 ± 4.4 cm/s.

We analyzed the correlation between the duration of monkey S’s decision and his movement speed at the single trial level in every of the 261 sessions he performed, and we found a significant correlation in 189 out of the 261 sessions. In each case of these 189 sessions with a significant correlation between decision duration and action speed, the longest decisions were followed by the fastest movements (Fig. S4, top left panel). We looked at the evolution of this relationship across sessions and, as we did for monkey G and monkey B in the main manuscript, we compared this evolution to the evolution of the duration of monkey G’s decisions in each of these sessions (Fig. S4, middle left panel). A visual inspection of these two metrics suggest a strong relationship, confirmed by a significant Pearson correlation analysis (r=0.36, p<0.0001, Fig. S4, bottom left panel). Thus, the strength of the positive correlation between decision duration and movement speed (the longest the decision, the fastest the movement) within trials through practice coincides with the increase of the duration of monkey S’s decisions through practice.

By analyzing the correlation between the duration of monkey Z’s decision and his movement speed at the single trial level, we found a significant correlation in 174 out of the 279 sessions. Among these 174 sessions with a significant correlation between decision duration and action speed, the longest decisions were followed by the fastest movements in 166 sessions (Fig. S4, top right panel). Finally, by assessing the relationship between this degree of coordination between decision duration and movement speed and the duration of decisions between sessions (Fig. S4, middle right panel), we found a significant correlation between the two parameters (Pearson correlation, r=0.13, p=0.026, Fig. S4, bottom right panel).

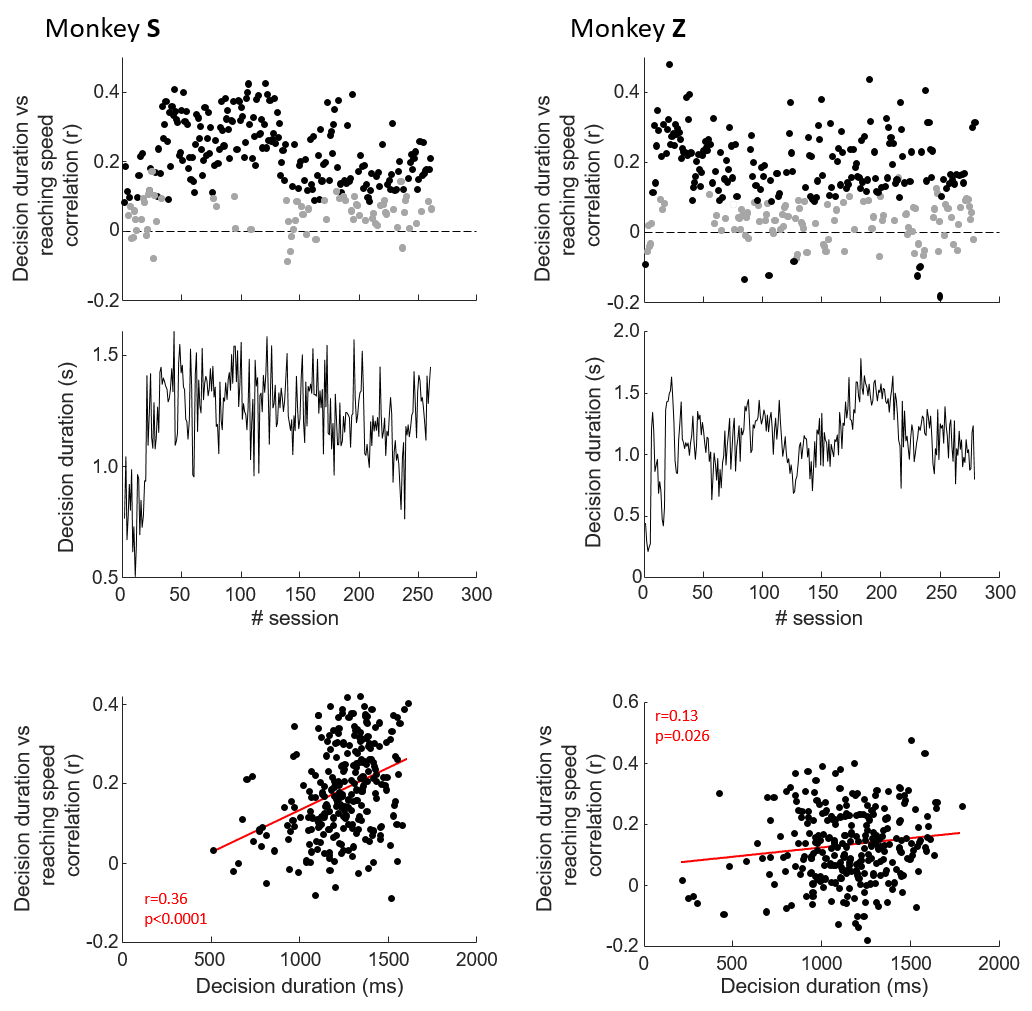

**Figure S4 - Relation between decision duration and arm movement speed at the single trial level in monkey S and monkey Z.** A - top panel: Pearson linear correlation between the duration of monkey S’s decisions (DD) and arm movement speed (velocity peak, VP) as a function of the session performed by the monkey. Each dot illustrates the value for a given session. A negative (positive) r value means that the longest the decision, the slowest (fastest) the movement. The black filled dots mark sessions in which the Pearson correlation is significant. Middle panel: Mean duration of monkey S’s decisions within sessions, as a function of the session performed by the monkey. Bottom panel: Relationship between the Pearson correlation between decision durations and the speed of monkey S’s movements and the mean duration of his decisions in each session. Each dot illustrates the value for a given session. The red line illustrates the result of a linear regression through the data. B – Same as in A for sessions performed by monkey Z.

Relationship between movement duration and movement peak speed

For both monkeys, with all trials collapsed across sessions, the relationship between movement duration and movement peak speed is highly significant (monkey G: n = 18954, Pearson correlation coefficient r = -0.26, p<0.0001; monkey B: n = 15630, r = -0.21, p<0.0001). The relationship is also significant (Pearson correlation, p < 0.05) at the session level in 89% of the 71 sessions performed by monkey G (96% if only correct movements are included, i.e. excluding inaccurate and too slow movements), and in 74% (80% for correct movements only) of the 46 sessions performed by monkey B. Finally, the correlation between movement duration and movement peak velocity is significant between sessions, for both monkeys, as can be seen in the figure below (Fig. S5).

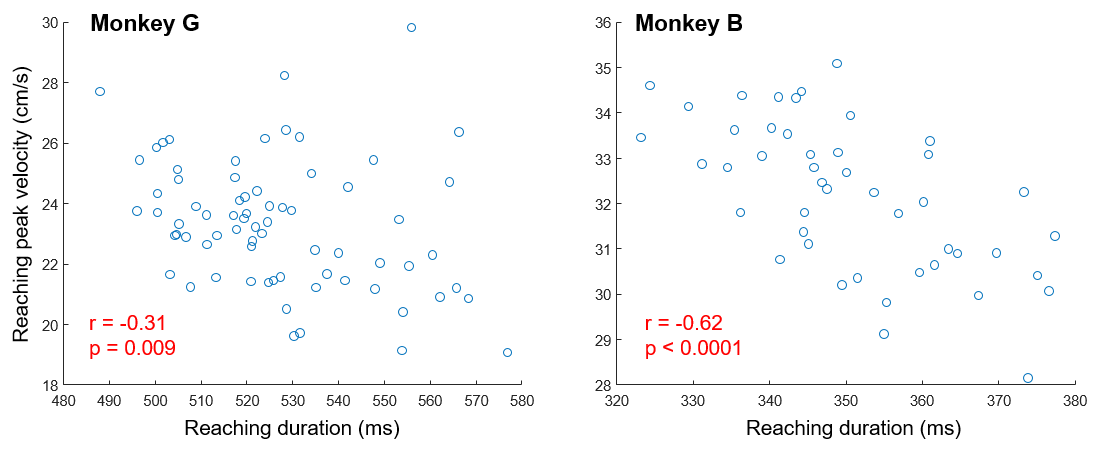

**Figure S5: Relation between the average reaching duration (abscissa) and the average reaching peak velocity (ordinate) in each session performed by monkey G and by monkey B.** Each blue dot illustrates data from a given session
